# The Visual Experience Evaluation Tool: A Myopia Research Instrument for Quantifying Visual Experience

**DOI:** 10.1101/2024.09.20.614212

**Authors:** David Sullivan, Aaron Nicholls, George Hatoun, Samuel Thompson, Cory Schwarzmiller, Fathollah Memarzanjany, Alyssa Gunderson, Alexander Danielson, John Lowes, Jon Petersen, Sam Backes, Micah Thomas, Hanna Eha, Lisa Rutherford

## Abstract

Current myopia research has demonstrated the role of extended visual experience in healthy ocular development. Optical cues and the spectrum, intensity, and temporal characteristics of light landing on the retina are all known factors affecting the development of the eye. However, there is still limited understanding as to which of these extrinsic factors are most important or how they interplay with intrinsic physical and neural differences between individuals. Part of the problem is inadequate tooling. Our team at Reality Labs Research created the Visual Environment Evaluation Tool (VEET), a non-commercial research instrument, to accelerate myopia research. In this paper, we describe the VEET’s physical design, sensor suite and capabilities, and the associated software which makes it well-suited for research of quantified visual experience.

## Introduction

Studies of myopia, both observational and experimental, have repeatedly suggested a link between visual experience and myopia onset and progression. Mandatory indoor education [1, 2], near work activities [3, 4], outdoor light exposure [5], and the intensity, spectral distribution and temporal characteristics of light near the eye [6] are all documented to affect ocular development.

Nevertheless, identifying the most consequential contributing factors has been challenging [7, 8]. Since ocular development continues until an individual is around 20-21 years of age, studies must be longitudinal and purely observational to respect participants’ privacy. Furthermore, existing research methodologies have severe limitations: Subjective reporting correlates poorly with objective measurements [9], and current-generation devices leave room for improvement in form factor, fidelity, and/or privacy.

The VEET addresses these challenges by integrating the highest quality spectral, illuminance, movement, and distance sensors into a robust, lightweight, all-day wearable form factor. To preserve the wearer’s privacy, there are no cameras or microphones: all the VEET measures is the light being reflected back from the environment. The resulting output can be aggregated across researchers, creating a time-sequenced dataset of light exposure measurements.

Improved research tools such as the VEET can enable researchers to better understand the impact of environmental inputs on myopia development and inform scientific models of eye growth.

### The Visual Environment Evaluation Tool

**Figure 1:**
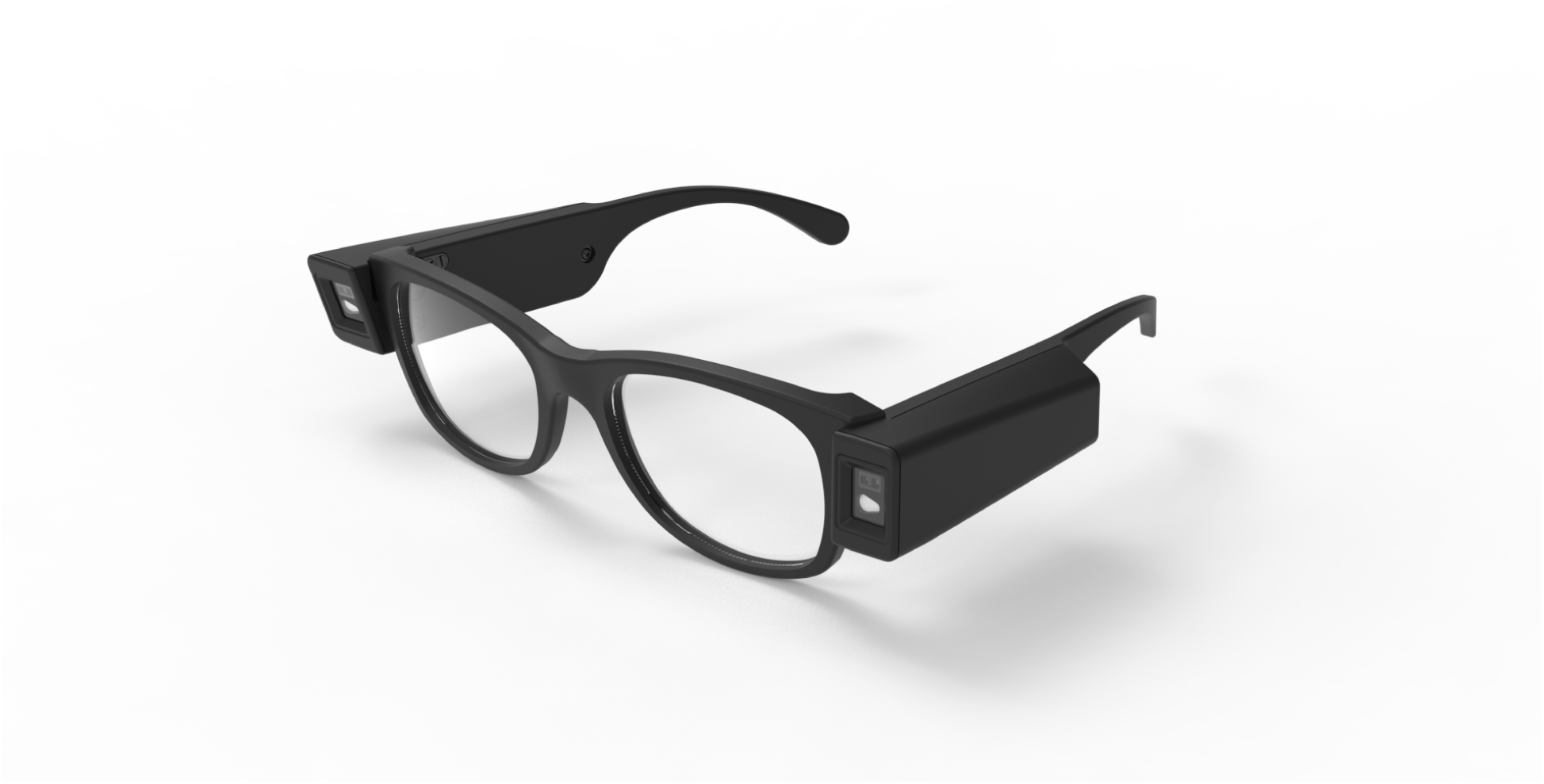
The VEET mounted on a commercially available frame.

The Visual Environment Evaluation Tool (VEET) is a pair of temple arms that gather quantitative data on characteristics of light near the eye without disrupting the wearer’s normal activities. Each temple arm is a standalone data-logging instrument that uses state-of-the-art sensors to measure photopic illuminance, visible spectrum, optical distance to near objects, and motion, all at sampling frequencies up to ½ hertz (every two seconds) for all day battery life. These robust solid-state sensors are industrially proven and commercially available, with excellent documentation and demonstrated long-term stability.

The VEET has no camera, no microphone, no bluetooth or wifi connectivity, and does not record any personally identifying information. The device logs data without compromising the wearer’s privacy or comfort.

Each sensor’s data output is fully exposed, allowing calculations to be confirmed or post-processed for specific purposes. The data is logged in a human-readable Comma Separated Values (CSV) file and is only accessible through a standard USB-C connection. The VEETManager, an open-sourced companion software application, allows live sensor viewing and device configuration and maintenance.

Each VEET temple arm comes with a battery that supports all-day use. With a recharge time of 2 hours on a standard USB-C wall charger, the battery allows for greater than 24 hours of continuous runtime at ½ Hz sensing rates.

If charged daily, the VEET has a data storage capacity of more than one year. Data is continuously logged at pre-configured rates whenever battery reserves are available or the device is charging on wall power.

### Form Factor and Fit

**Figure 2:**
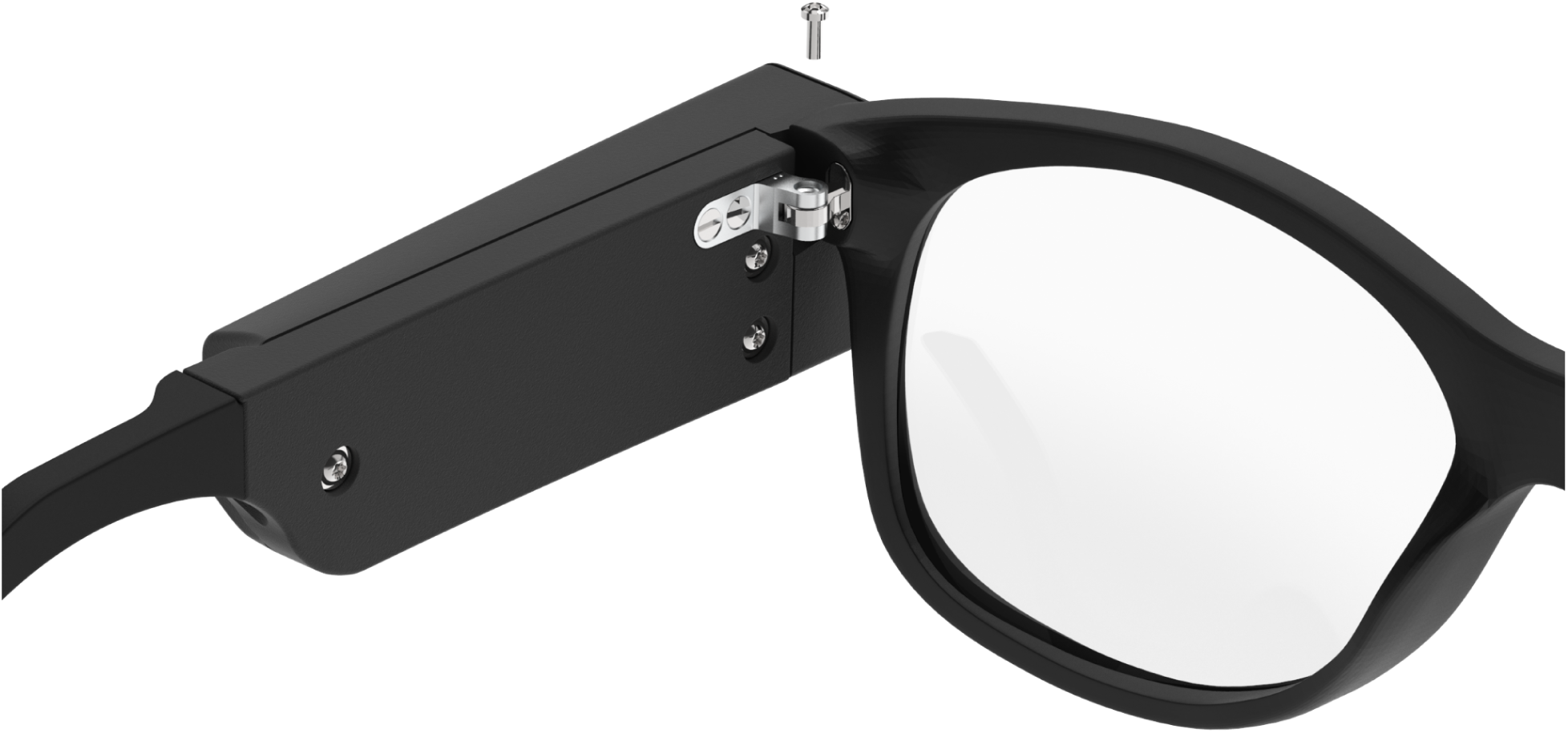
VEET installation on glasses frames.

The VEET ships with a customization kit, enabling compatibility with a wide range of commercially available glasses frames. Researchers select from four different hinges and four exchangeable arm extensions (125mm-145mm), which allows for finding the best fit for each participant. To assemble, the researcher simply replaces the existing temple arms with the VEET temple arms and reuses the original hinge screws.

Each temple arm has a mass of approximately 20 grams, light enough for all-day wear. While the arms could be used separately, we recommend using them as a pair to maintain proper balance.

### Electronics and Sensing Layout

**Figure 3:**
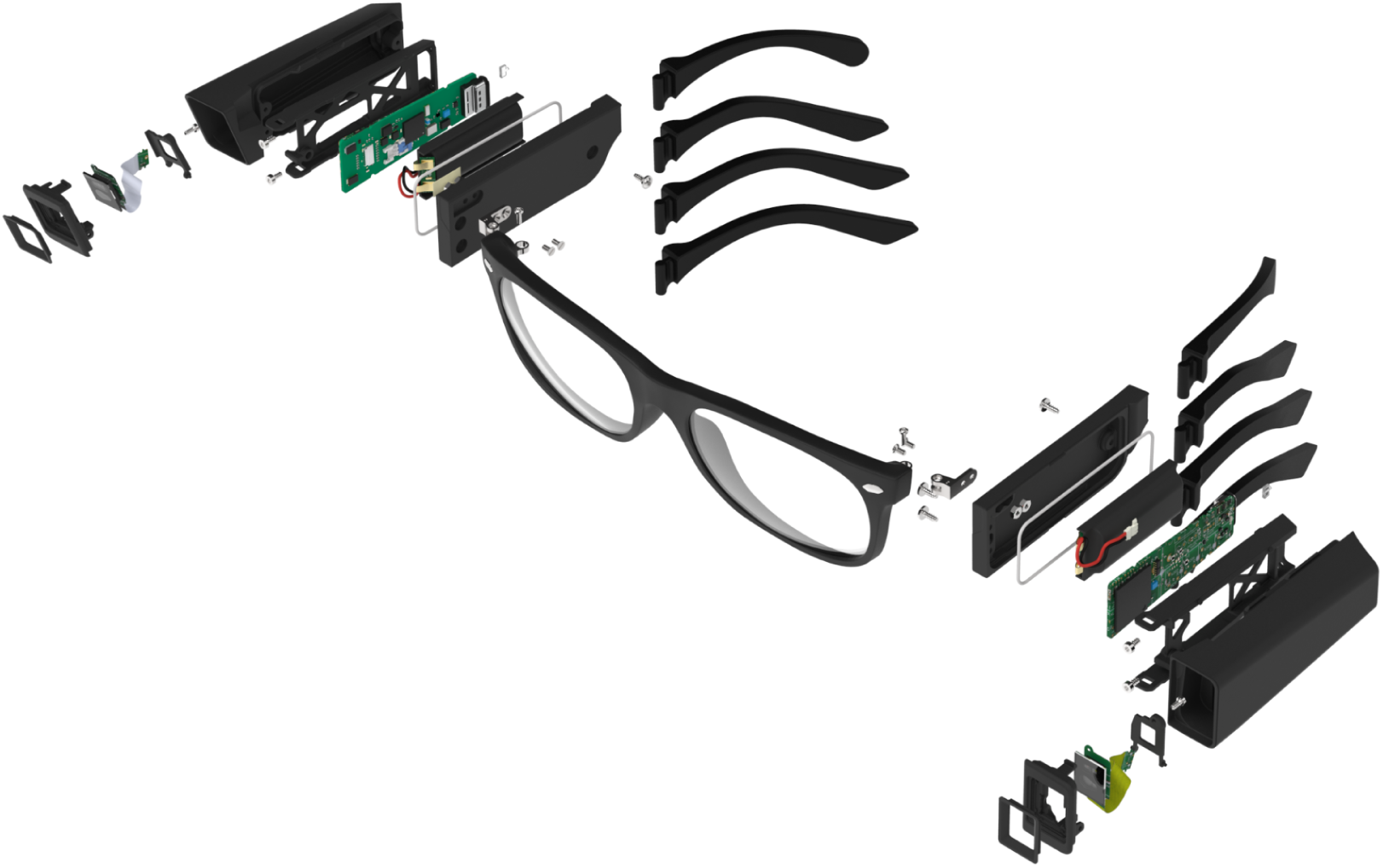
The internal components of the VEET.

The VEET enables all-day sensing at relatively high rates while minimizing bulk volume and weight. Each temple arm contains an identical electronics package, consisting of three forward facing sensors:

● Time of Flight or Range detection (ams OSRAM part number TMF8828)
● 11 Channel Multi-Spectral meter (ams OSRAM part number AS7341)
● Ambient Light Sensor (ams OSRAM part number TSL2585)

The forward-facing sensors are internally connected to the power, control and communications systems, including the following key components:

● One Inertial Measurement Unit (Bosch part number BMI270)
● Microchip ARM Cortex-M7 microcontroller
● 350 mAh lithium polymer battery
● Minimum 16gb flash memory storage
● USB Type-C connection

To cover a range of potential gaze behaviors, the right temple arm’s forward-facing sensors are aimed 20 degrees downward and 4 degrees towards the sagittal plane while the left temple arm’s sensors are perpendicular to the glasses frame. The Inertial Measurement Unit (IMU) on both temple arms is housed along the temple in identical orientations.

**Figure 4:**
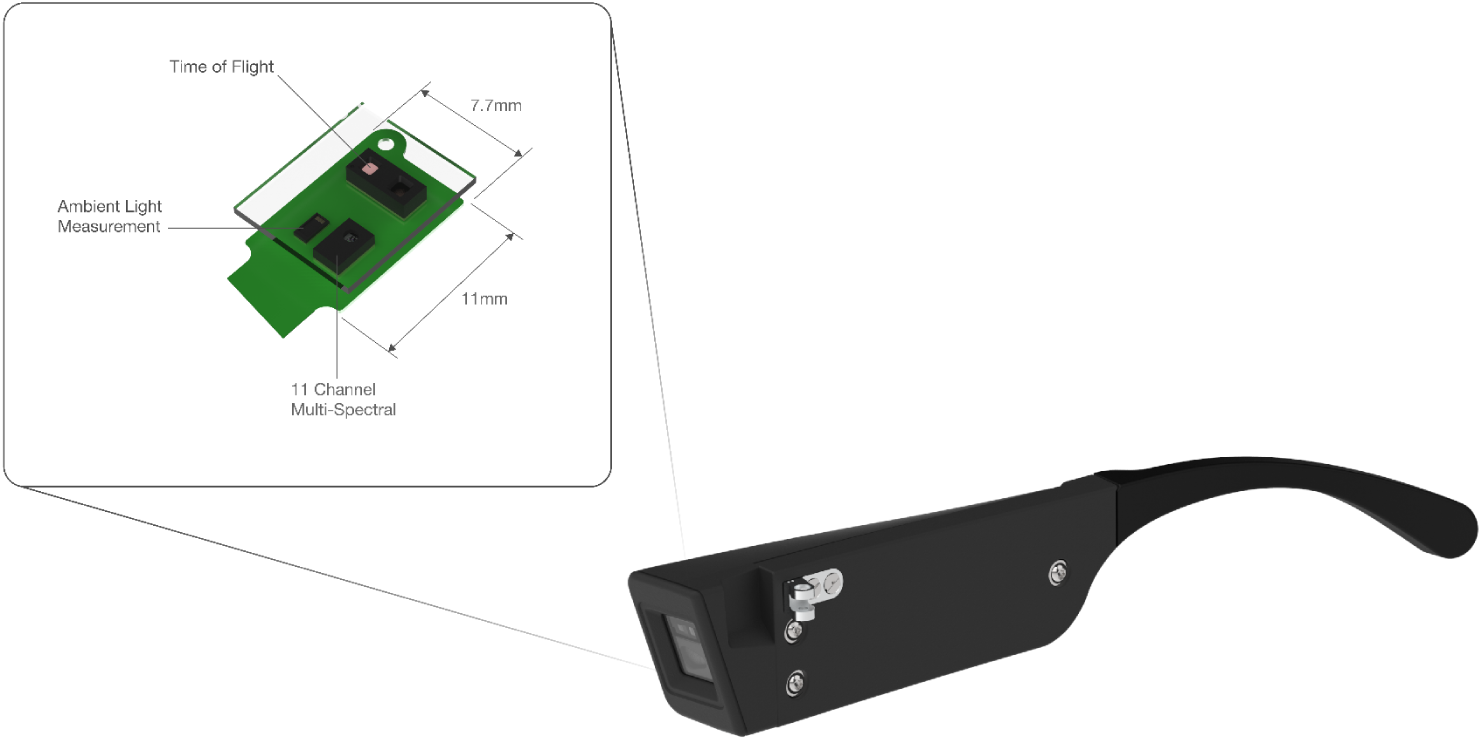
VEET front-facing sensor layout.

### Optical Configuration

The three forward-facing sensors are packaged behind a custom optical stackup that allows all three sensors to be on the same circuit board. The stackup starts with a 0.55mm thick AGC Dragontrail coverglass—an aluminosilicate glass with excellent scratch and impact resistance—commonly used across many consumer and automotive touchscreen applications. The glass has a nearly flat light transmission curve that exceeds 90% in the visible and infrared ranges of interest (400-1200 nm). Below 400 nm, the transmission becomes more variable and falls off to less than 60% by 300 nm. Ultraviolet wavelengths below 400 nm will have responsivities that deviate from the manufacturer’s datasheet-specified response curve [10,11]. The optical sensors packaged in the VEET have a specified full spectral responsivity from near 300 nm to approximately 1150 nm. Ultraviolet sensitivity is impacted by this optical configuration. Calculations for illuminance and spectral response incorporate accommodations to account for the falloff in sensitivity in this region.

To minimize packaging volume, the field of view of each sensor is reduced from full cosine response. The aperture geometry is determined by the tallest component—the Time of Flight Sensor—and the overall housing width. For the Spectral and Ambient Light Sensors, a 0.13mm thick lambertian diffuser is required to evenly distribute light from the environment onto the sensors and to eliminate bright spots. The diffuser is a Kimoto 100PU with double-sided diffusion coating, a haze of 89.5%, and a transmission of 66%. It is placed between a compliant rubber boot and the top aperture piece, and is retained into the housing by a sealed bezel. This geometry results in apertures with an effective minimum field of view as shown in Figure 5.

**Figure 5:**
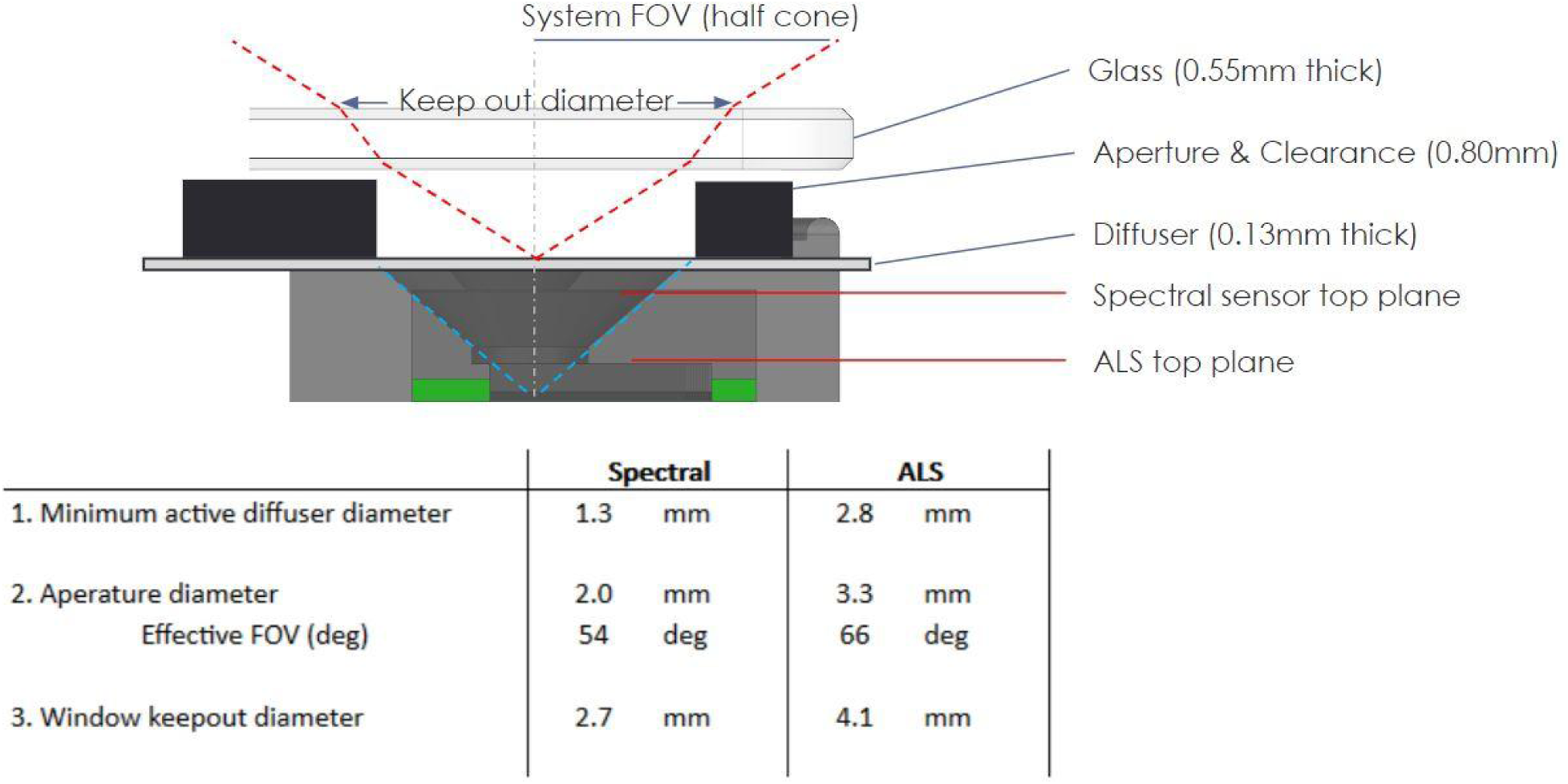
VEET optical stackup assembly.

## Sensor Details

### Ambient Light Sensor (ams-OSRAM TSL2585)

The Ambient Light Sensor’s primary function is to provide photometric data directly in lux. It also includes broad spectrum (UVA, photopic visible, and infrared) counts and light flicker detection.

The raw photopic response of the sensor closely matches the human eye’s response with the following exceptions: The sensor has some responsiveness in the infrared light regime, and the specific optical components in the light path influence the light measurement, requiring an assumption of the light source type to predict an accurate lux value. If sources of light that contain large amounts of infrared radiation, such as direct outdoor sunlight or incandescent bulbs, are not accounted for, they will negatively influence the lux calculation.

A Labsphere Spectra UT-1000-S Polychromator was used to evaluate the responsivity of the spectral sensor in the visual domain as packaged in the VEET. As shown in the Normalized Spectral Responsivity chart in Figure 6, the ALS sensor’s visible wavelength response closely matches the sensor manufacturer’s datasheet [10] with a slight shift in peak wavelength towards the ultraviolet.

**Figure 6:**
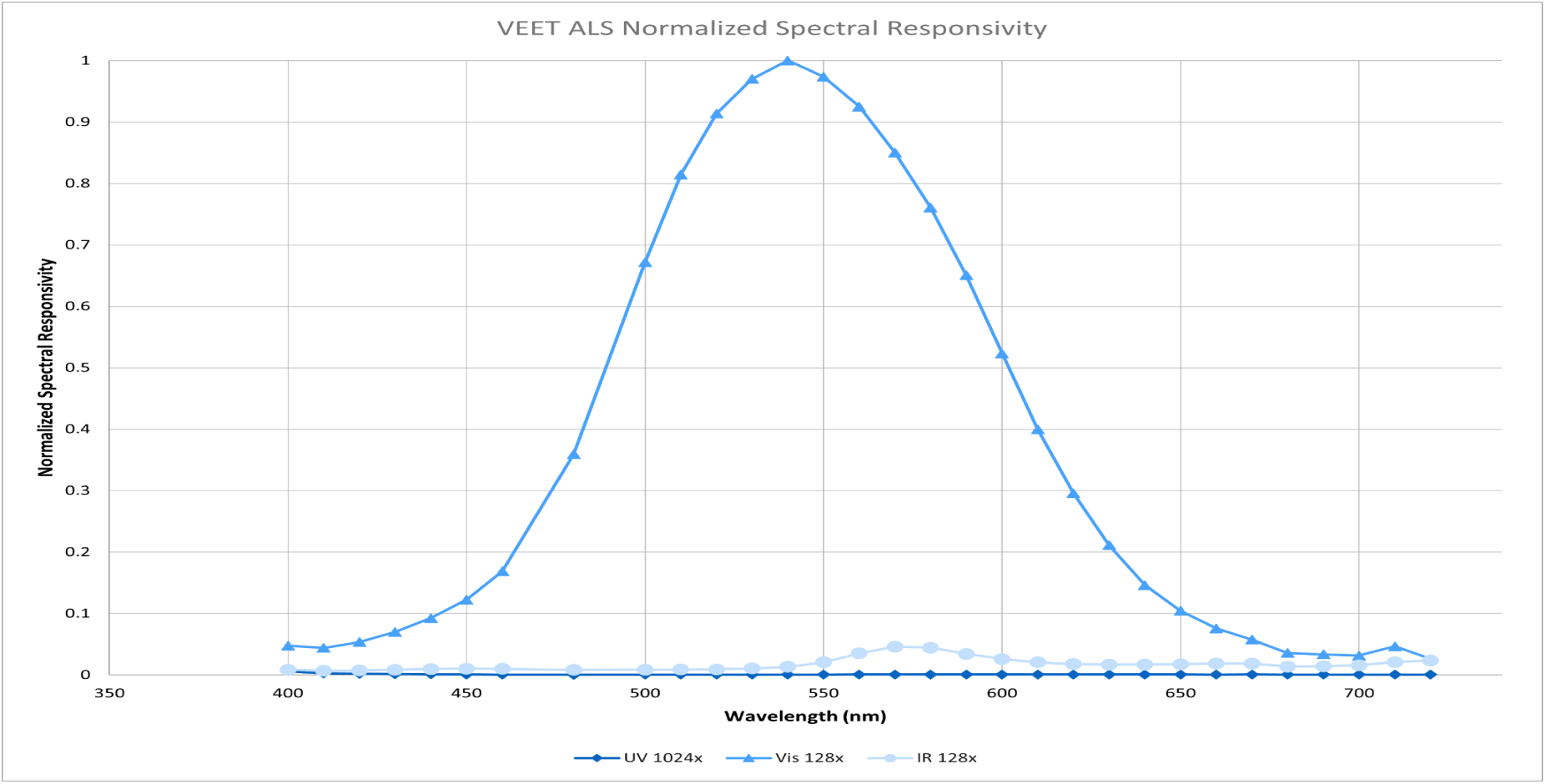
ALS normalized spectral responsibility.

To ensure the VEET can predict lux accurately in both low IR and high IR lighting environments, the VEET was exposed to a wide variety of common lighting sources and mixed environments and compared to a calibrated Minolta CL500a Illuminance Spectrophotometer to allow compensation of the raw sensor output response. As shown in Figure 7, the resulting compensation takes the form of a four piecewise equation that accurately predicts lux values ranging from typical low IR source illumination (e.g. LED lighting) to typical high IR source illumination (e.g. various outdoor lighting conditions and incandescent sources). The firmware compares the infrared, ultraviolet, and visible responses and adjusts the lux calculation for every sample based on these experimentally determined coefficients. Exposing the VEET to lighting sources not typically found in the living environment may result in inaccurate lux values being recorded.

**Figure 7:**
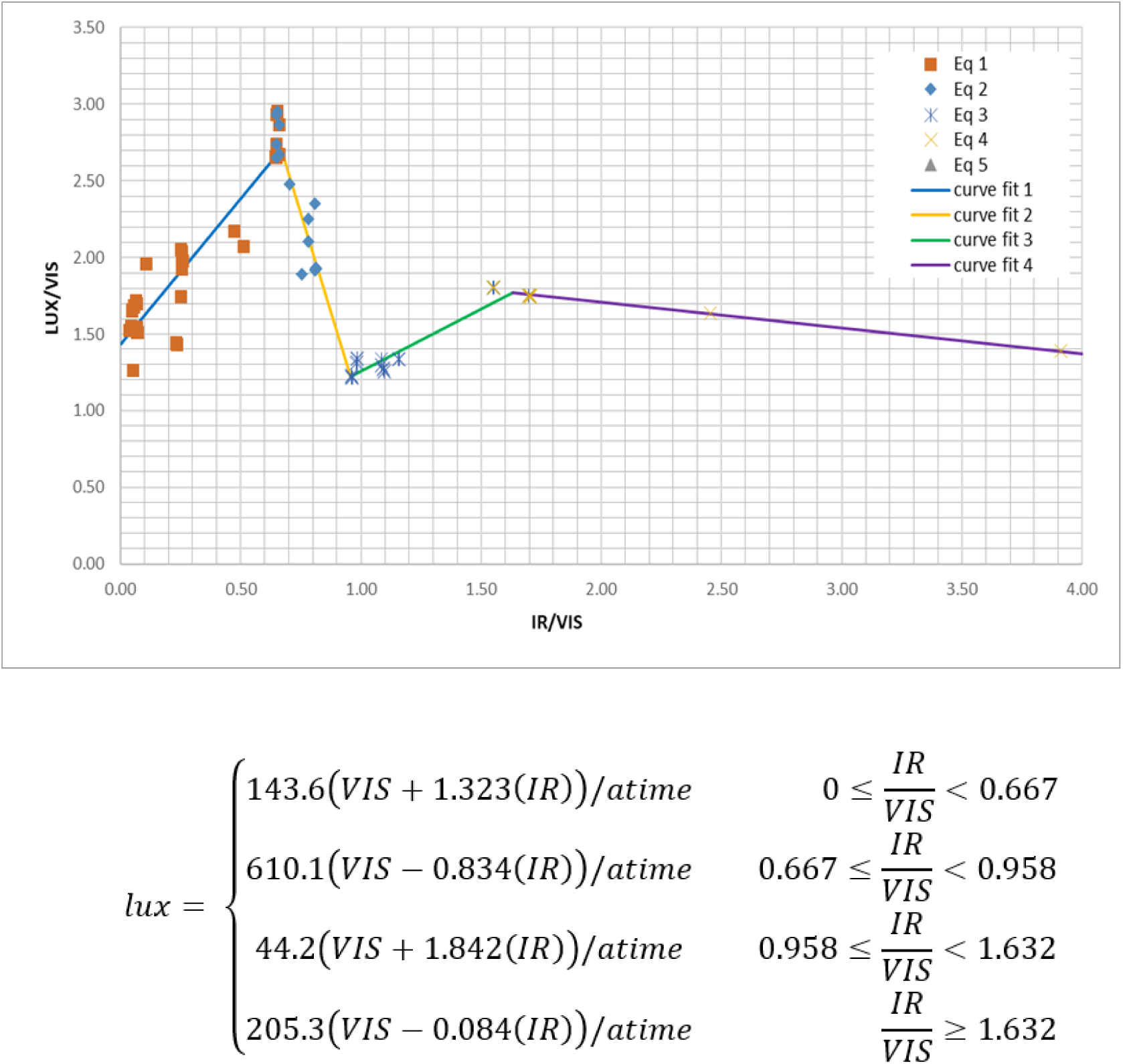
Plot of recorded data and curve fits used to generate the piecewise lux equation.

The VEET has been validated to produce reasonable lux values from approximately 0.1 lux to >100,000 lux. Observed performance is better than +/−10% accuracy, as compared to a high accuracy spectrophotometer, except at very low light levels (less than 0.5 lux), where the VEET may overestimate lux by up to 15% with the current lux calculation method.

As shown in Figure 8, during all-day, continuous sampling, the VEET may report illuminance values below 0.1 lux in dark environments. Accuracy below 0.1 lux has not been thoroughly evaluated due to limitations with existing validation instrumentation. Improvements in very low light accuracy may be feasible with modifications to the lux calculation methods and further testing. Each sensor’s data output is fully exposed: calculations can be confirmed, or post processed for specific purposes, enabling updating of existing data sets with new approaches.

**Figure 8:**
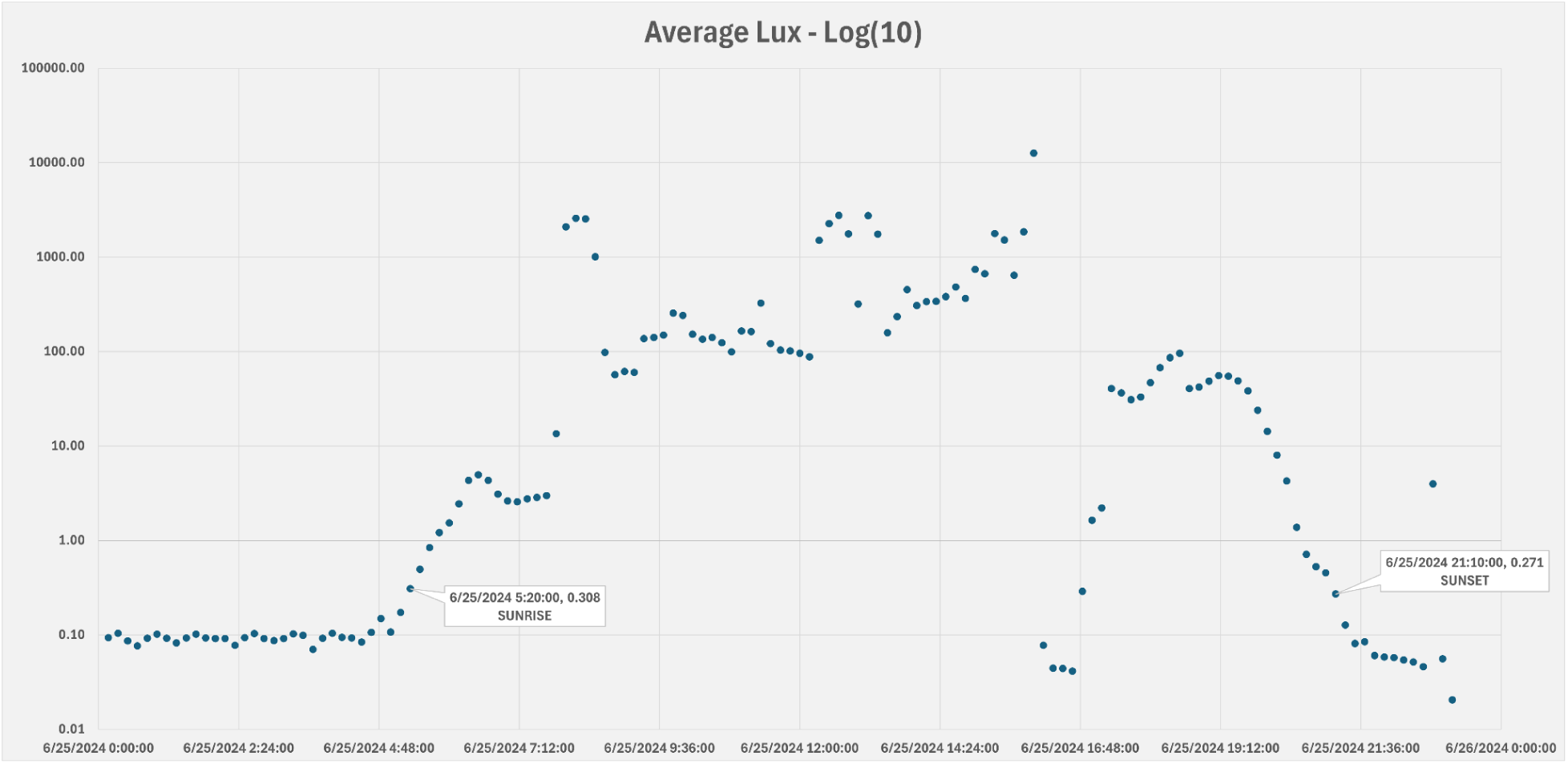
A typical day’s lux recording.

### Angular Response

As shown in Figure 9, the Ambient Light Sensor has an aperture-limited angular response. While the sensor itself has a near cosine response, packaging in the VEET housing reduces the field of view to 66 degrees. Light that exceeds this effective field of view is not received by the sensor.

**Figure 9:**
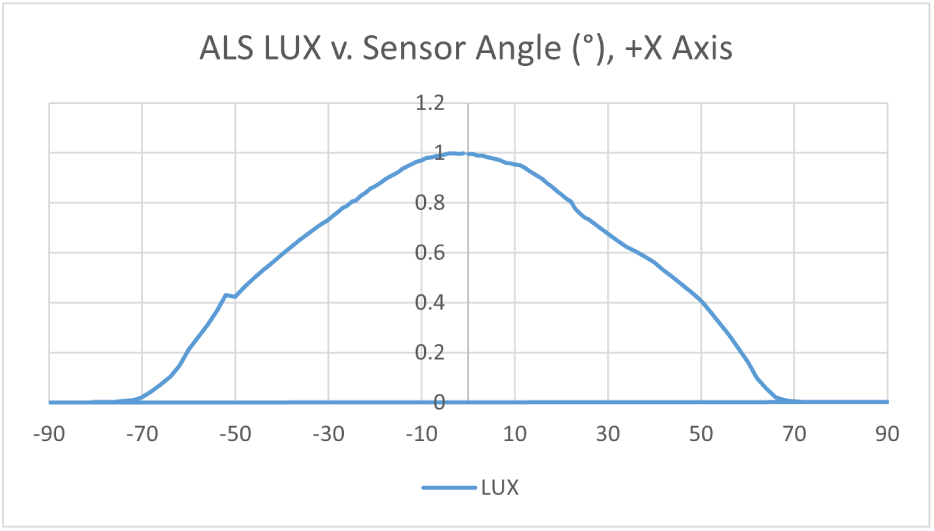
Typical angular response of the VEET’s Ambient Light Sensor.

### Flicker Detection

The Ambient Light Sensor natively samples the light environment at 14 kHz for the duration of the prescribed integration (typically 100ms), detecting flicker rates up to 7kHz. Performance has not been thoroughly validated in complex flicker environments, but is reasonable when exposed to typical office lighting (60 or 120Hz) or screens (120 or 240 Hz), providing secondary validation of an indoor or otherwise artificially lit environment.

### Spectral Sensor (ams-Osram AS7341)

The Spectral Sensor is an 11-channel multi-spectral sensor for color detection and spectral analysis. As shown in the Measured Spectral Responsivity chart in the sensor manufacturer’s datasheet [11], the Spectral Sensor uses 8 narrow channels plus 3 broad channels to measure the amount of light in the environment. The spectral response is defined in wavelengths from approximately 350 nm to 1000 nm. A Labsphere Spectra UT-1000-S Polychromator was used to evaluate the responsivity of the spectral sensor in the visual domain as packaged in the VEET. Figure 10 shows the typical responsivity differs slightly from the manufacturers datasheet, with the VEET having slightly higher relative sensitivity at wavelengths below 555 nm. These differences are characterized for each device at manufacturing and incorporated into the individual calibration.

**Figure 10:**
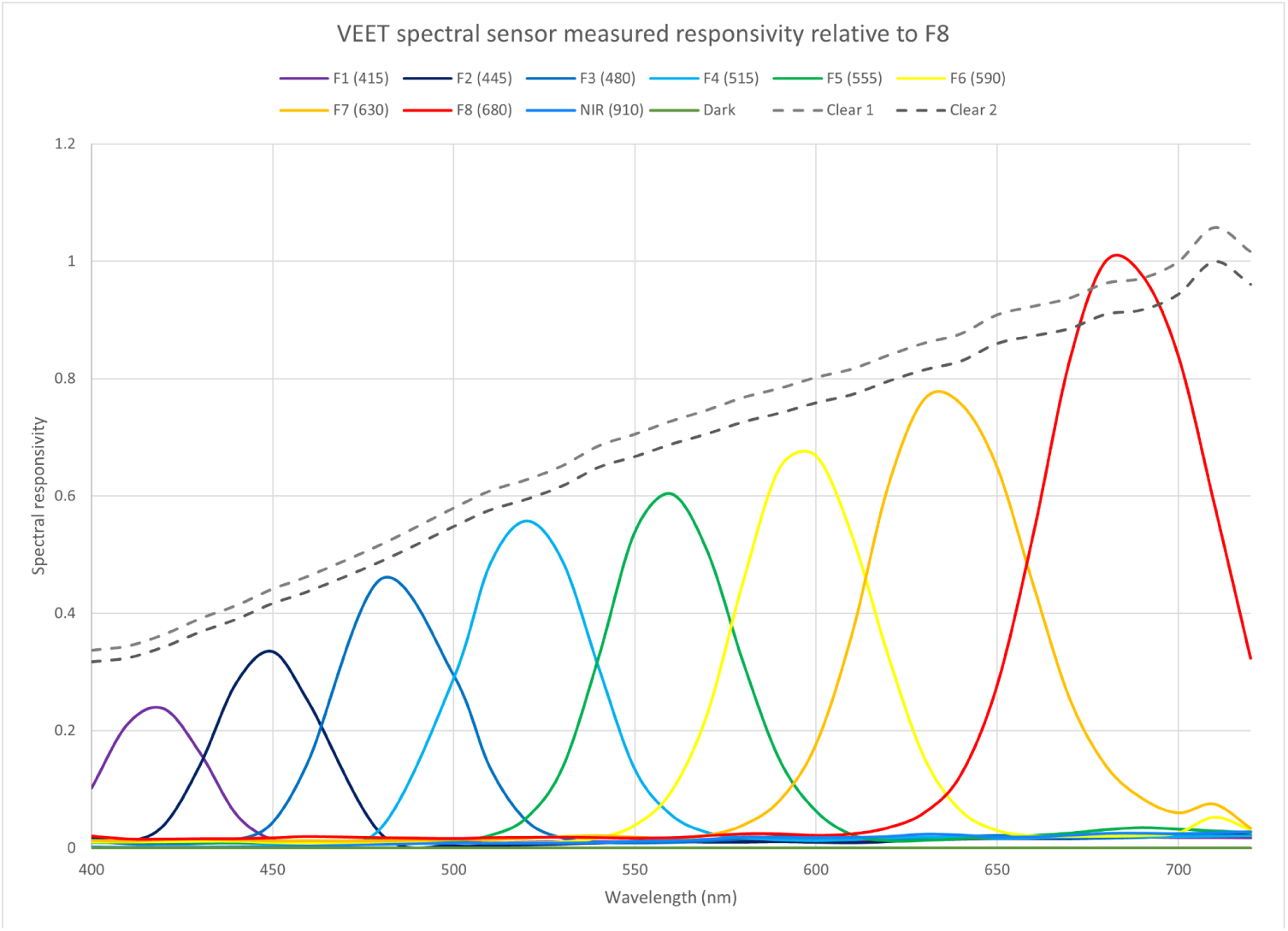
Spectral Sensor normalized spectral responsibility.

Each channel of the Spectral Sensor is a photodiode that produces and records raw counts in its region of sensitivity, as well as information on noise level in the environment, as shown in Table 1. The VEET sets the Integration Time and Gain for the sensor and records it with the sensing data.

**Figure 11:**
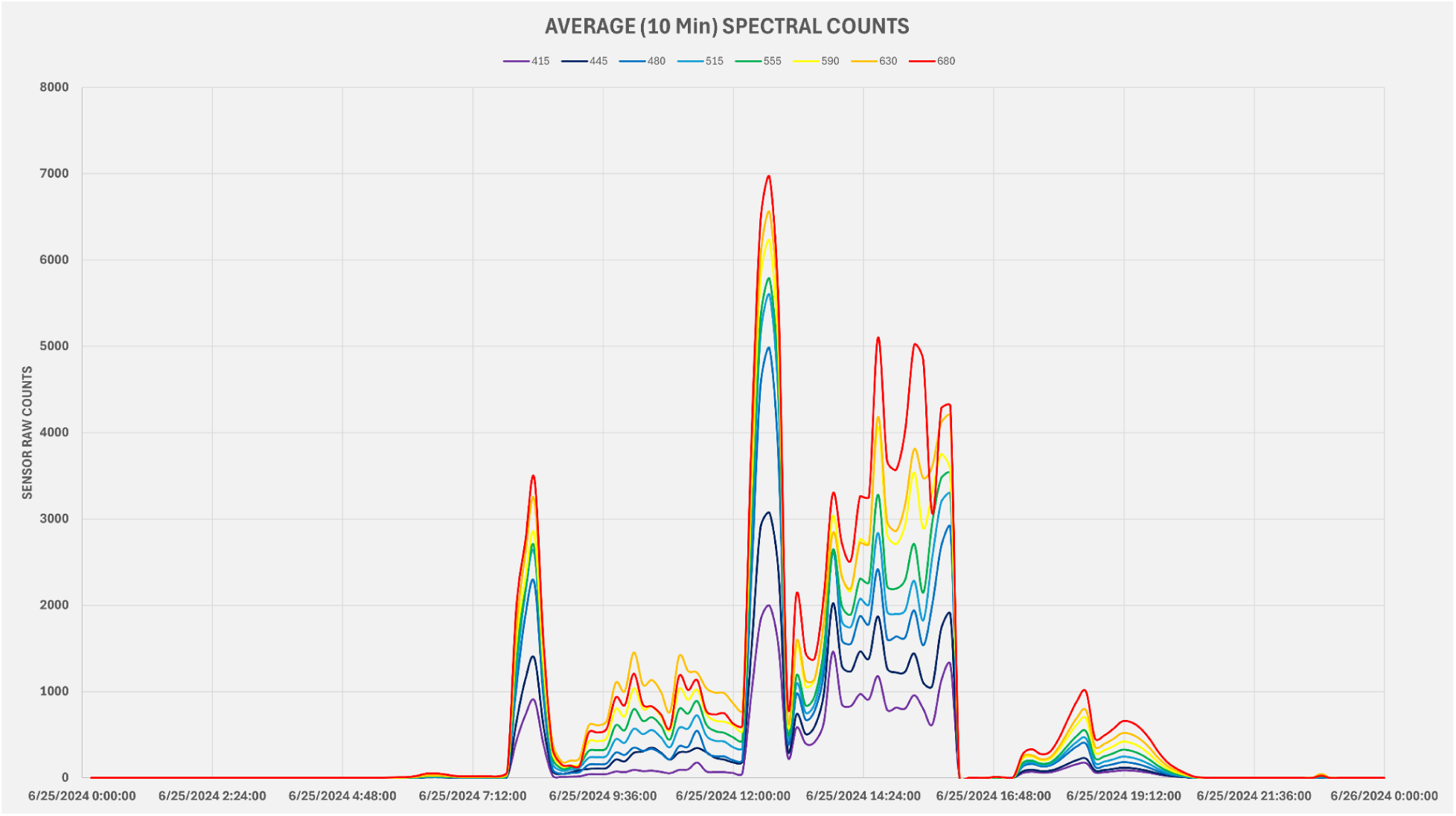
A typical day’s Spectral recording.

**Table 1:**
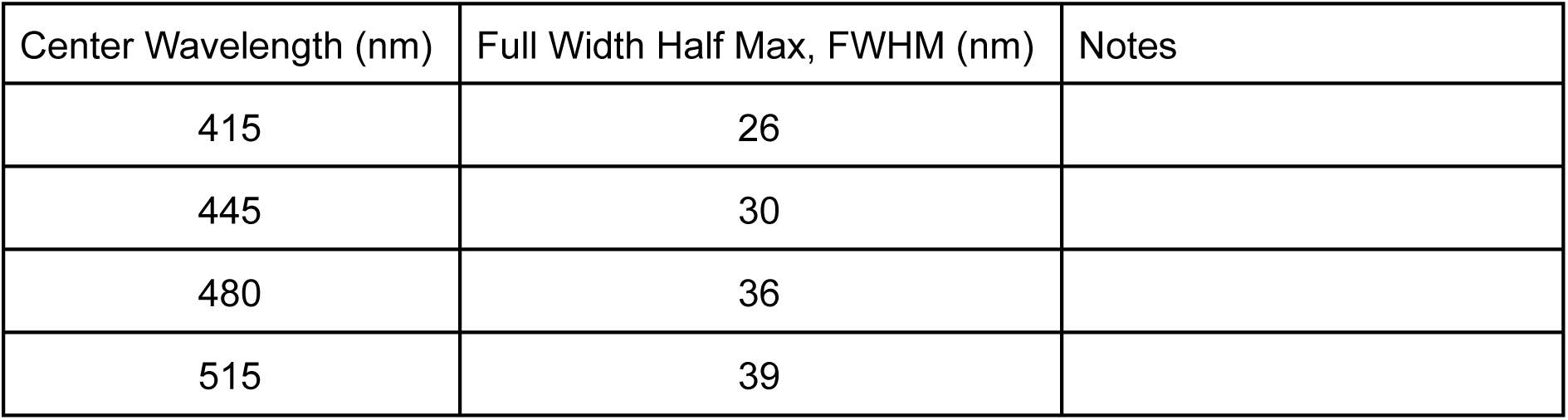
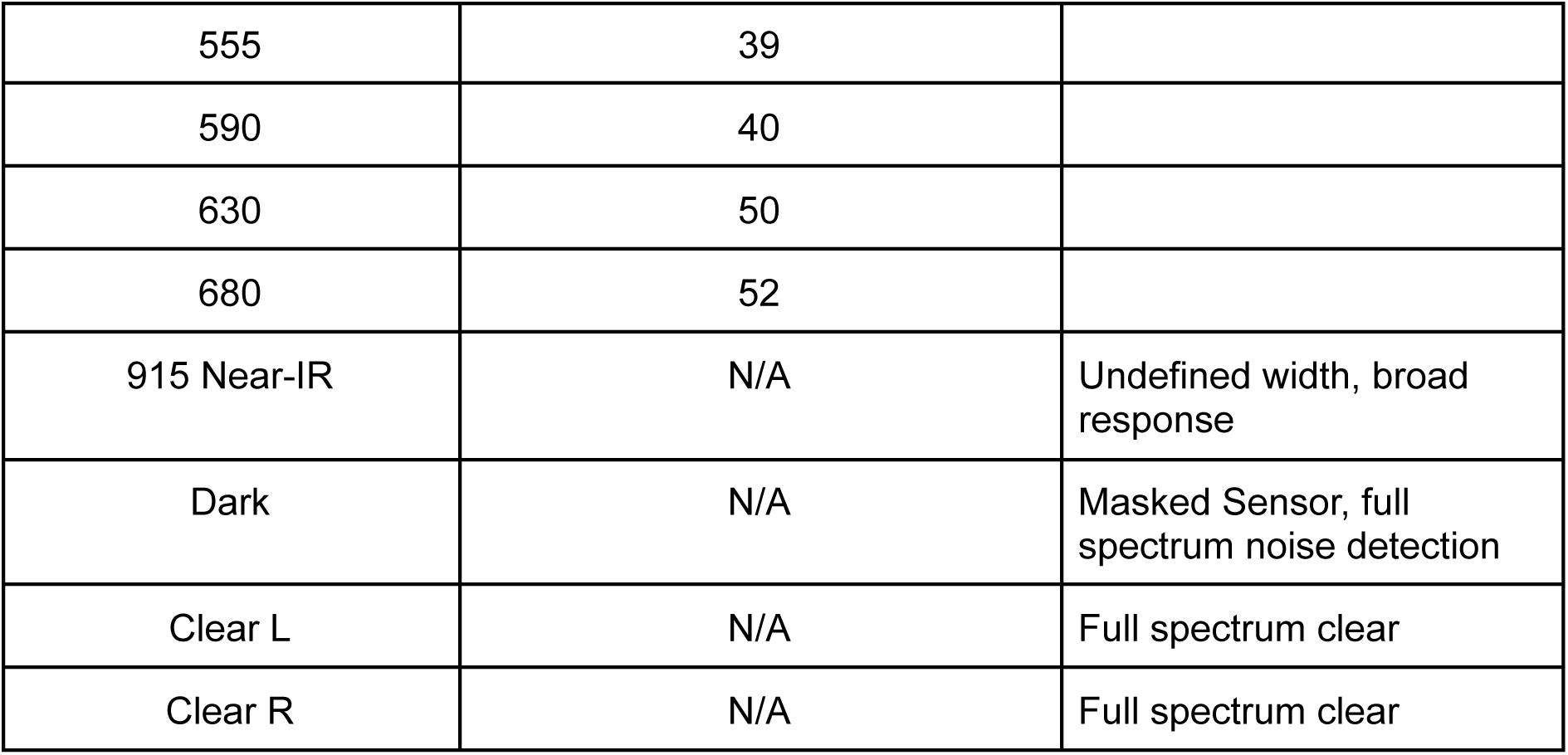
AS7341 Optical Channel Summary.

As packaged and configured in the VEET, the sensor provides spectral information at light levels down to approximately 5 lux.

This data is sufficient to be post-processed to predict correlated color temperature (CCT) and/or spectral power distribution (SPD). There are multiple methods of varying complexity to estimate the SPD and CCT based on the raw counts recorded by this sensor. It is beyond the scope of this paper to document these approaches.

### Angular Response

As shown in Figure 12, the Spectral Light Sensor has an aperture limited angular response. While the sensor itself has a near cosine response, the VEET housing reduces the field of view to half angles of approximately 60 degrees horizontally and 50 degrees vertically. The effective field of view is asymmetric due to the mechanical packaging of the sensor with slightly asymmetrical diffuser implementation.

**Figure 12:**
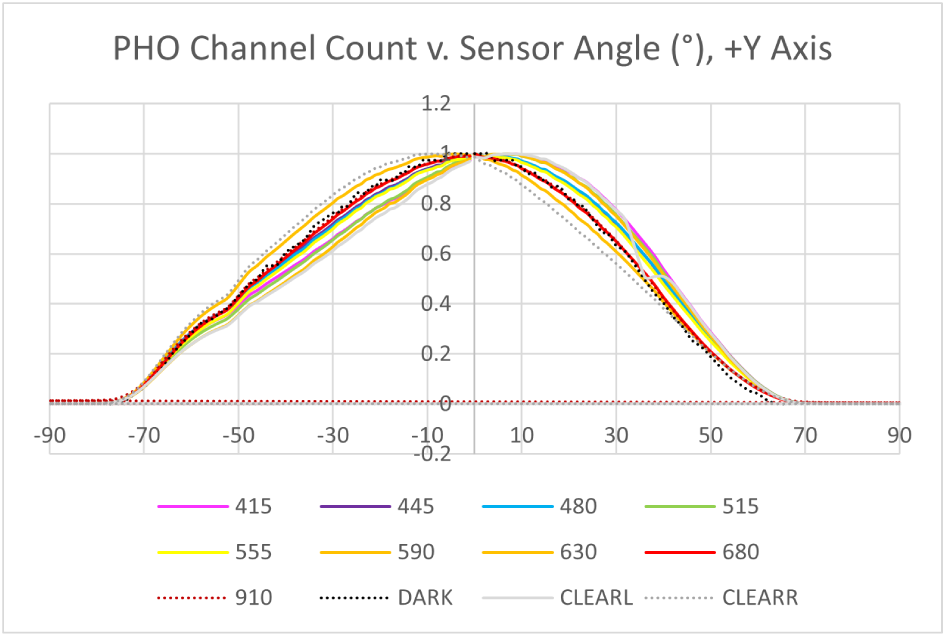
Angular response of the Spectral Light Sensor when packaged in the VEET.

### Time of Flight Sensor (ams-OSRAM TMF8828)

**Figure 13:**
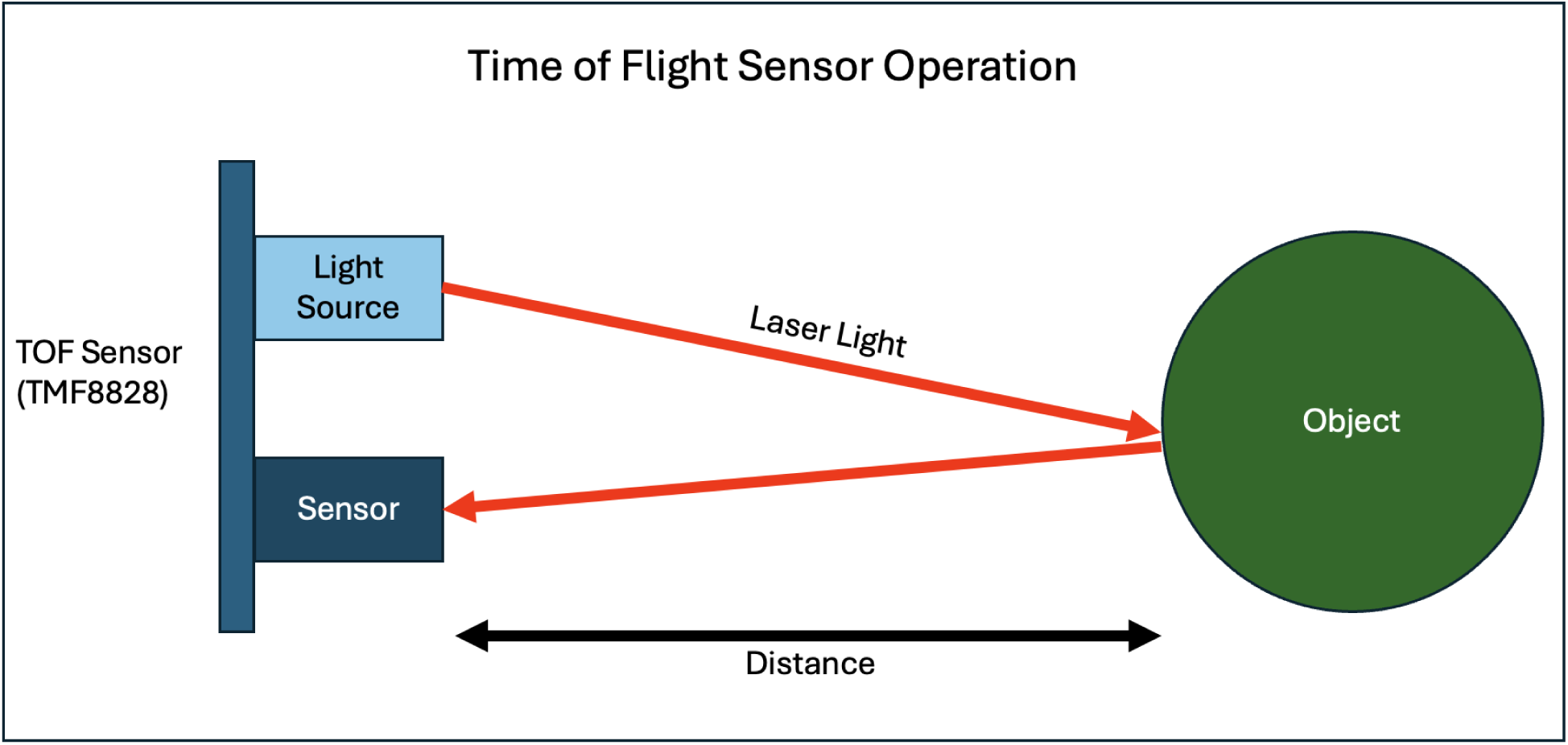
Simplified diagram of Time of Flight sensing.

The VEET’s TMF8828 ranging sensor is a direct time-of-flight (dToF) module capable of multizone depth measurements at a maximum FoV of 41° x 52° deg (approximately –26 to +26 degrees vertically and –20.5 to +20.5 degrees horizontally) and a resolution of 8 x 8 measurement zones.

The Time of Flight Sensor (ToF) illuminates the nearfield environment with an infrared light utilizing a class 1, eye safe, Vertical-Cavity Surface-Emitting Laser (VCSEL) and measures the time it takes for the light to return from any reflected objects. Figures 14 and 15 illustrate the same days’ data visualized as a time series and as distribution of sensed distances.

**Figure 14:**
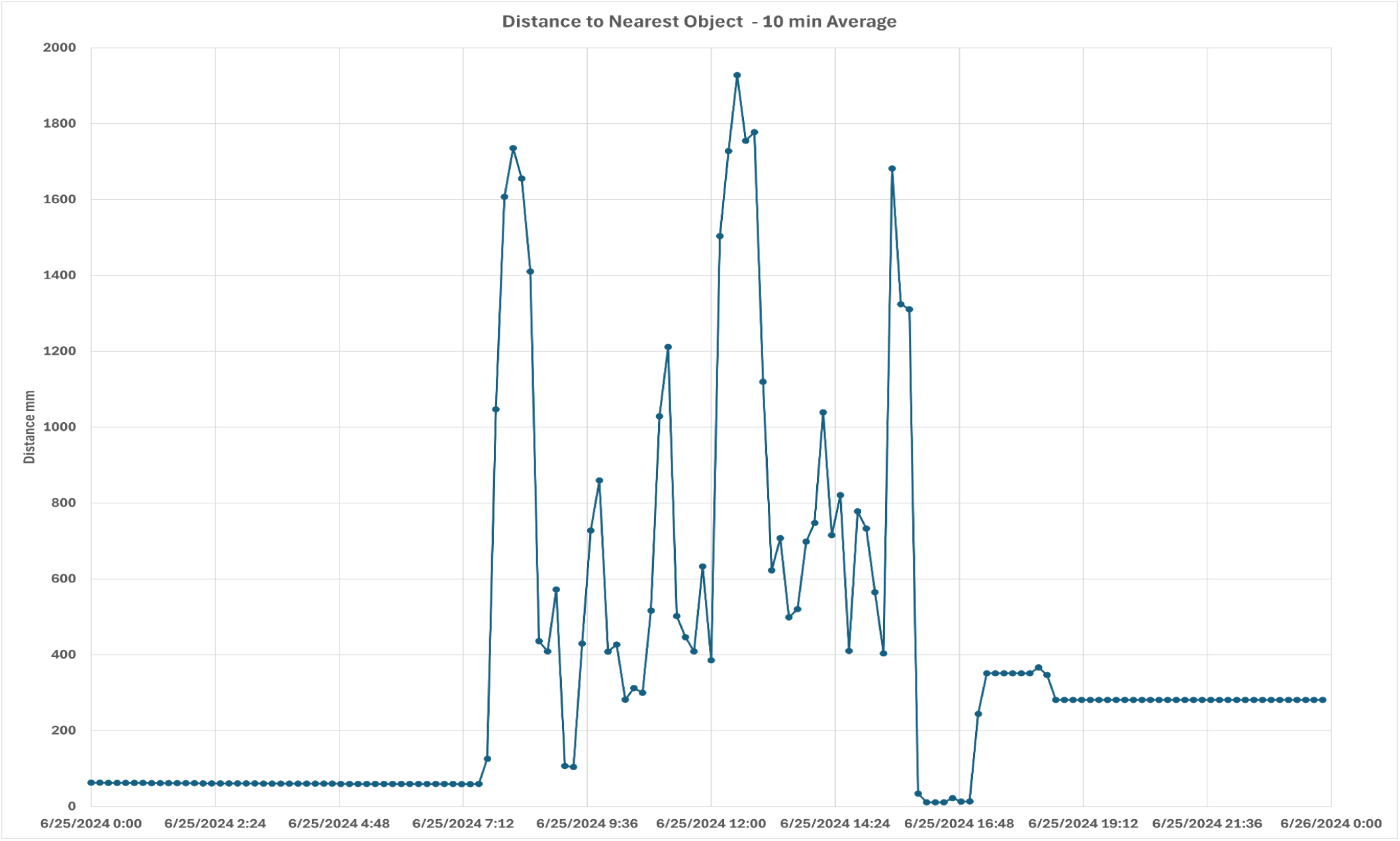
A typical day’s Time of Flight sensing data, aggregated at 10 minute intervals.

**Figure 15:**
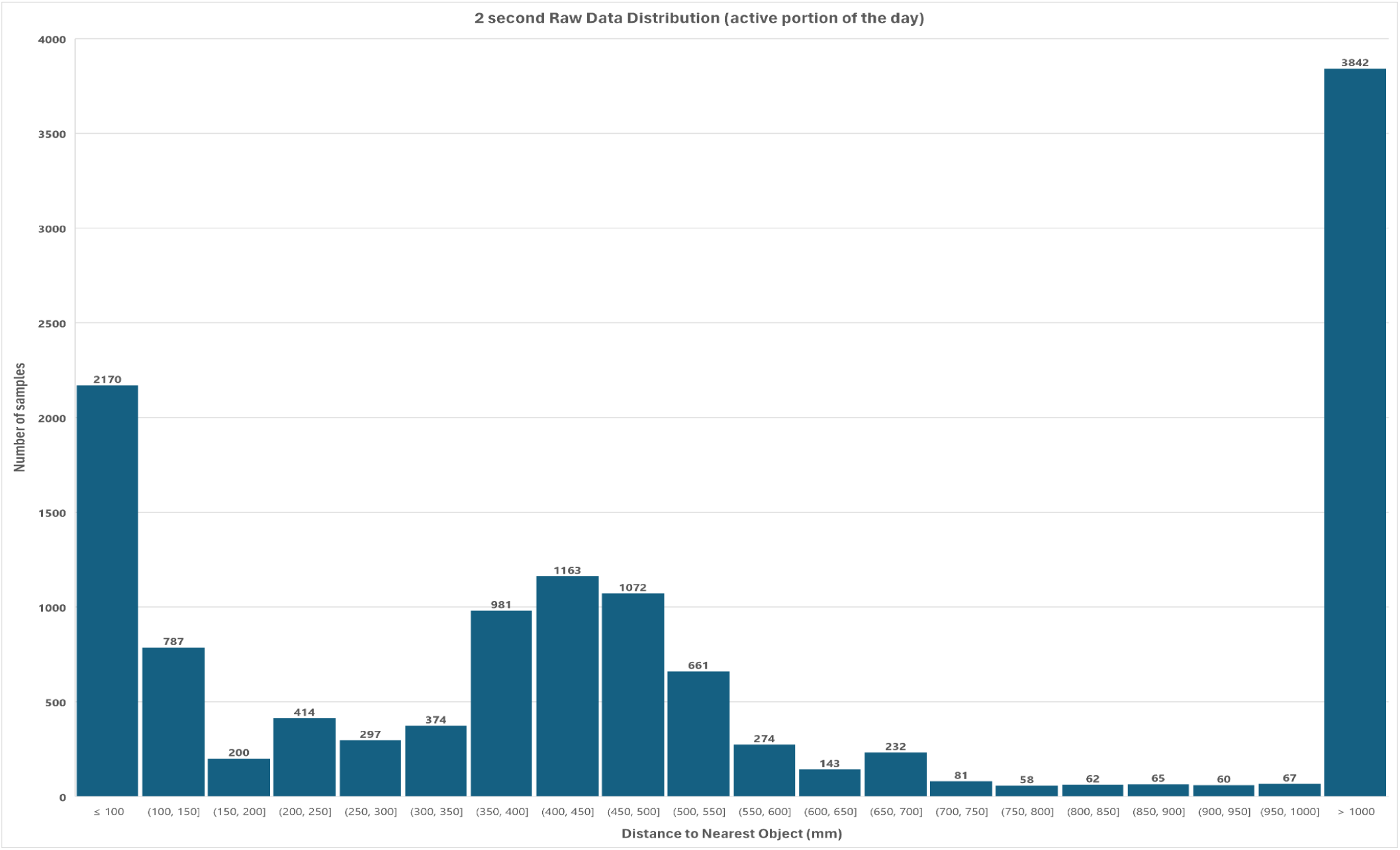
Distribution of a typical day’s Time of Flight sensing data.

To cover a range of potential gaze behaviors, the right temple arm’s sensors are aimed 20 degrees downward and 4 degrees towards the sagittal plane while the left temple arm’s sensors are perpendicular to the glasses frame. This feature of the VEET requires careful comparison of the data between devices as the same scene may produce significantly different distance results.

### Operation

During a ranging operation, the VCSEL pulses tens of thousands of times to illuminate the scene. Each specified zone collects these photons and creates a histogram of photon count against time. The following is an example histogram plot for a TMF8828 operating in 3×3 mode, for a total of 9 zone measurements per ranging operation [12].

**Figure 16:**
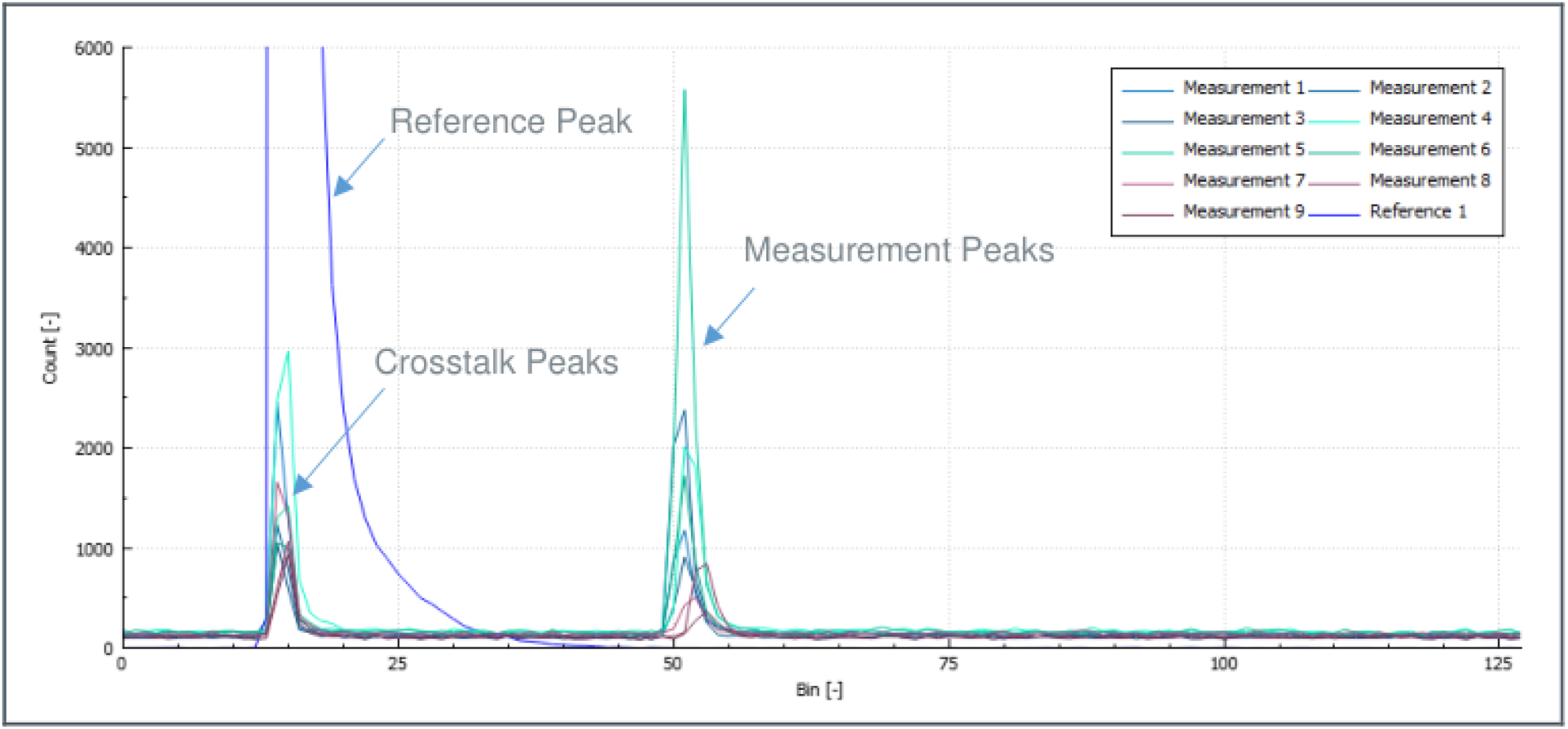
Example Time of Flight sensor histogram output, 3 x 3 mode.

The VEET captures four values for each of the 64 sensing zones: two values each (the Distance and Confidence) for up to two objects detected.

Distance (mm): The distance of a detected object, measured by the peaks of the photon count.

Confidence (0-256): A relative correlation to the width of the photon count histogram, Unitless.

The VEET records a total of 128 pairs of values for Distance and Confidence each time the sensor is sampled. If the Time of Flight Sensor detects no object in a zone, the Distance and Confidence values for that zone are 0. Minimum sensing distance is approximately 10mm for a fully occluded sensor.

The value of the Distance for a zone is the true distance from the Time of Flight Sensor and is not corrected to a flat plane. The sensor will report that the center of the field of view is closer than the edges when observing a flat plane. Simply averaging all 64 data points may slightly overestimate the shortest distance to near objects when measuring controlled scenes. Analysis of dToF data in complex, real-world, visual environments requires curation based on the research question being asked.

### Validation

To validate the ToF Sensor, we developed a fixture to accurately step the VEET along a controlled linear axis in 50mm increments, measuring the distance to a 300mm x 300mm flat plane test target with the ability to control obliqueness (see Figure 17).

**Figure 17:**
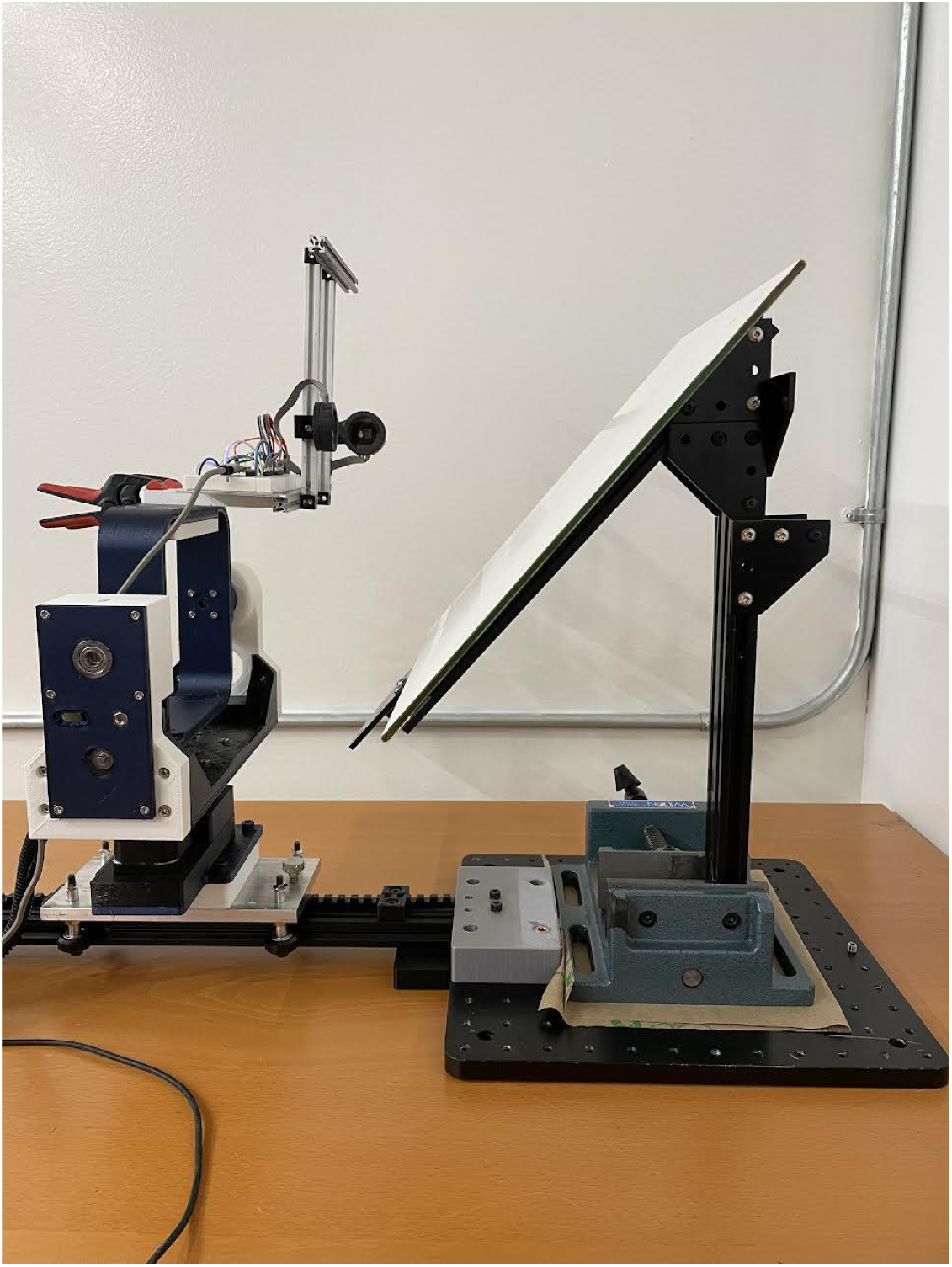
Fixture used to validate the VEET’s ToF Sensor.

The Time of Flight Sensor has a specified +/3% of full scale accuracy [12]. Our testing largely showed that the device performed within these tolerance bounds and had better accuracy in the middle sensing zones as compared to the edge zones. Furthermore, our testing shows that a confidence threshold of 30 (out of 255) was a reasonable cutoff confidence in a controlled environment. See typical center and edge responses in Figures 18 and 19 below.

**Figure 18:**
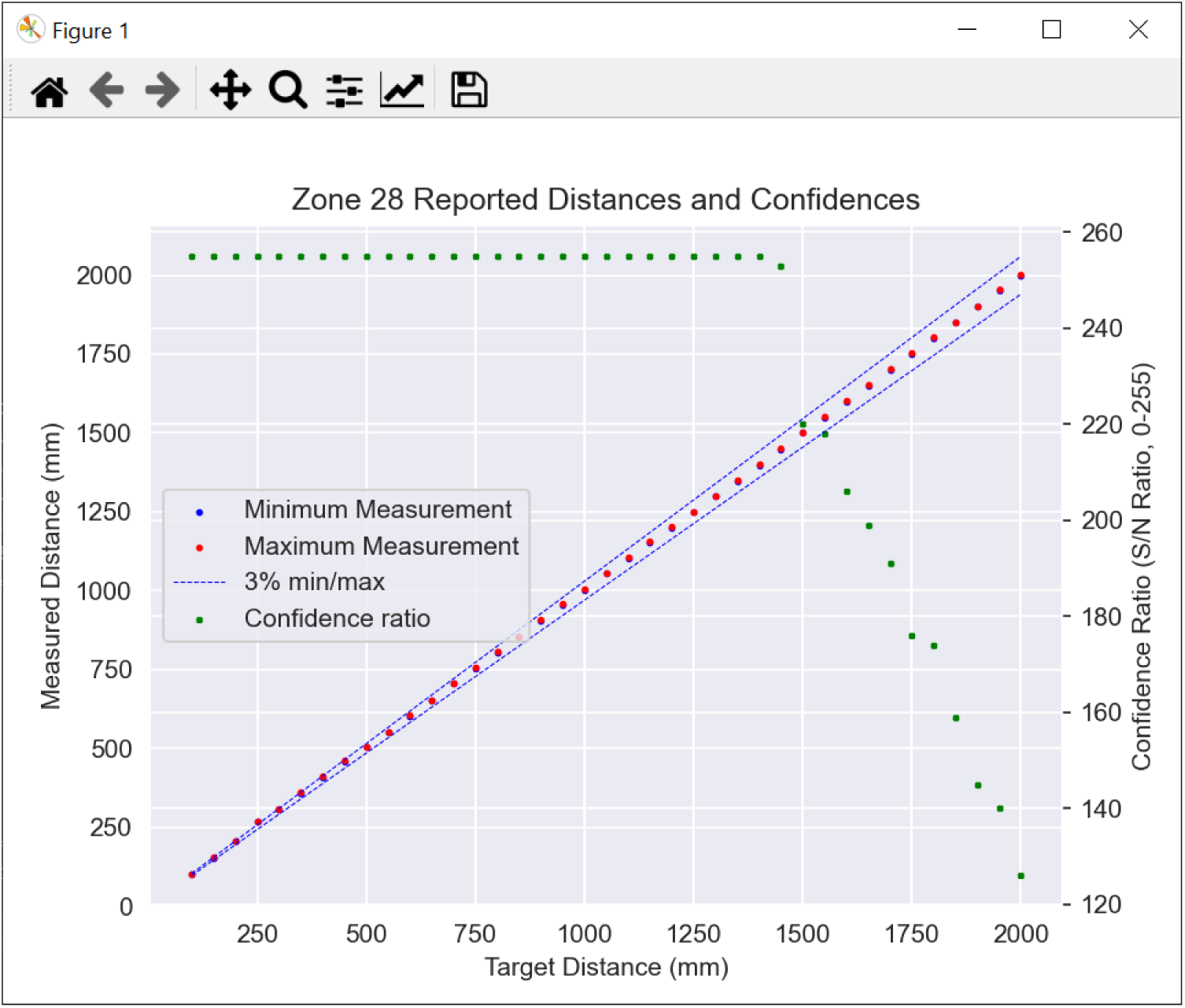
Center zone TOF response 50-2000mm.

**Figure 19:**
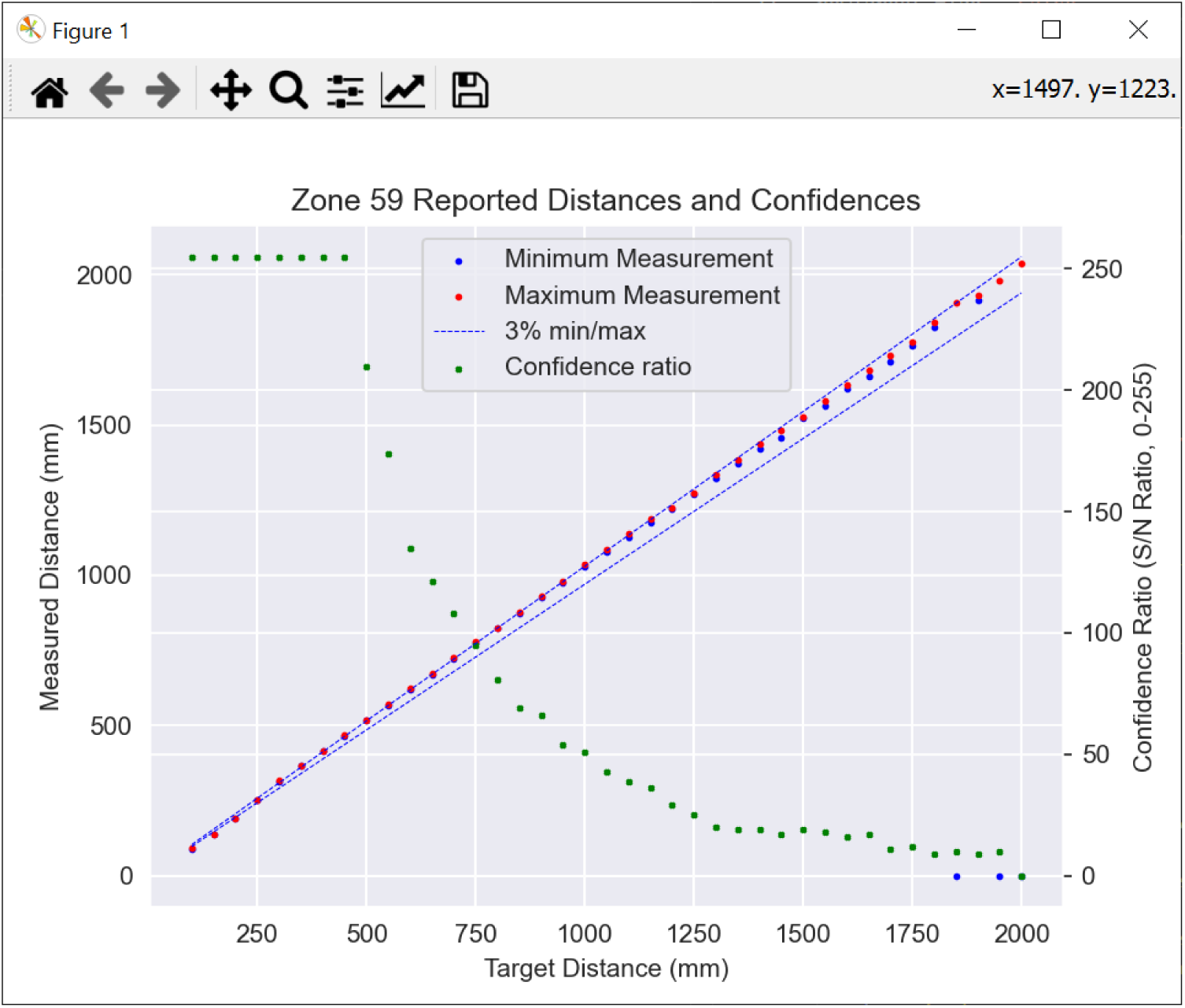
Edge zone TOF response 50-2000mm.

Multiple surfaces and obliqueness angles were evaluated. Based on these results, no calibration of the device is required.

### Environmental influences

Since the device utilizes an infrared VCSEL to illuminate the environment, lighting that has high levels of IR or objects with low IR reflectivity can both negatively affect the range and confidence levels reported. In highly controlled testing environments the manufacturer claims up to 5000 mm of distance measurement capability. In our testing 3000 mm is feasible as a reliable center distance upper limit, but is rarely seen in real world device testing. Our experience suggests typical indoor distance detection is reliable in a worn device up to 2000 mm. In bright, full sunlight environments the max range detection may drop to as low as 1000 mm.

Transparent objects can alter the distance measurement or appropriate interpretation of the results. For example, auto glass may reflect the IR signal and reasonably show the distance to the glass, even though the user is not typically focused on that object. Testing of individual scenarios is required if the VEET needs to detect objects through a visibly transparent object.

### Inertial Measurement Unit (Bosch BMI270)

The Inertial Measurement Unit consists of a three-axis accelerometer and a three-axis gyroscope that combine to provide information about movement and orientation. The sensor also provides temperature as a secondary output.

**Figure 20:**
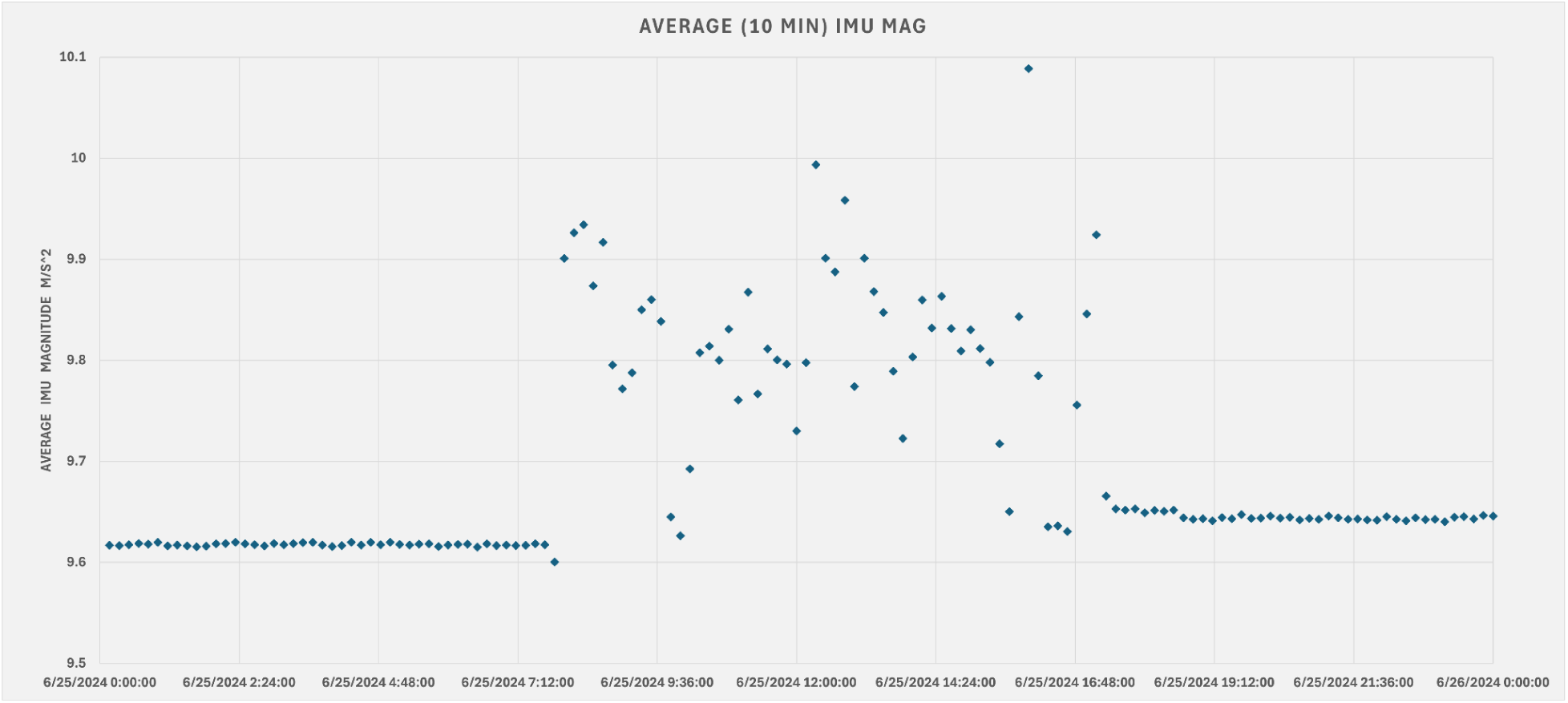
Distribution of a typical day’s Internal Measurement Unit sensing data.

The IMUs are housed in the side of each temple arm and are mounted in the same orientation. They do not need to be adjusted for the pointing angle of the other sensors packaged in the VEET. The IMU records acceleration and angular velocity on the device at the moment it is sampled.

Acceleration in the X, Y, and Z directions are recorded directly in meters/second^2 with a measurement range of +/−4g (+/−39.2 m/s^2), a sensitivity error up to +/−0.4%, and a zero-g (resting) offset up to .2 m/s^2 of total magnitude.

Rotation rates about the X, Y, and Z axis are recorded directly in Degrees per Second (dps) with a measurement range of +/−2000 degrees per second, a sensitivity error up to +/−0.4%, and an observed zero-rate offset up to +/−2 dps.

The IMU additionally stores the internal temperature in degrees celsius. This temperature is recording circuit board temperatures and has not been evaluated for correlation to ambient temperature.

No additional calibration is performed during production, as the manufacturer’s specified implementation is sufficient to support the VEET’s purposes.

**Figure 21:**
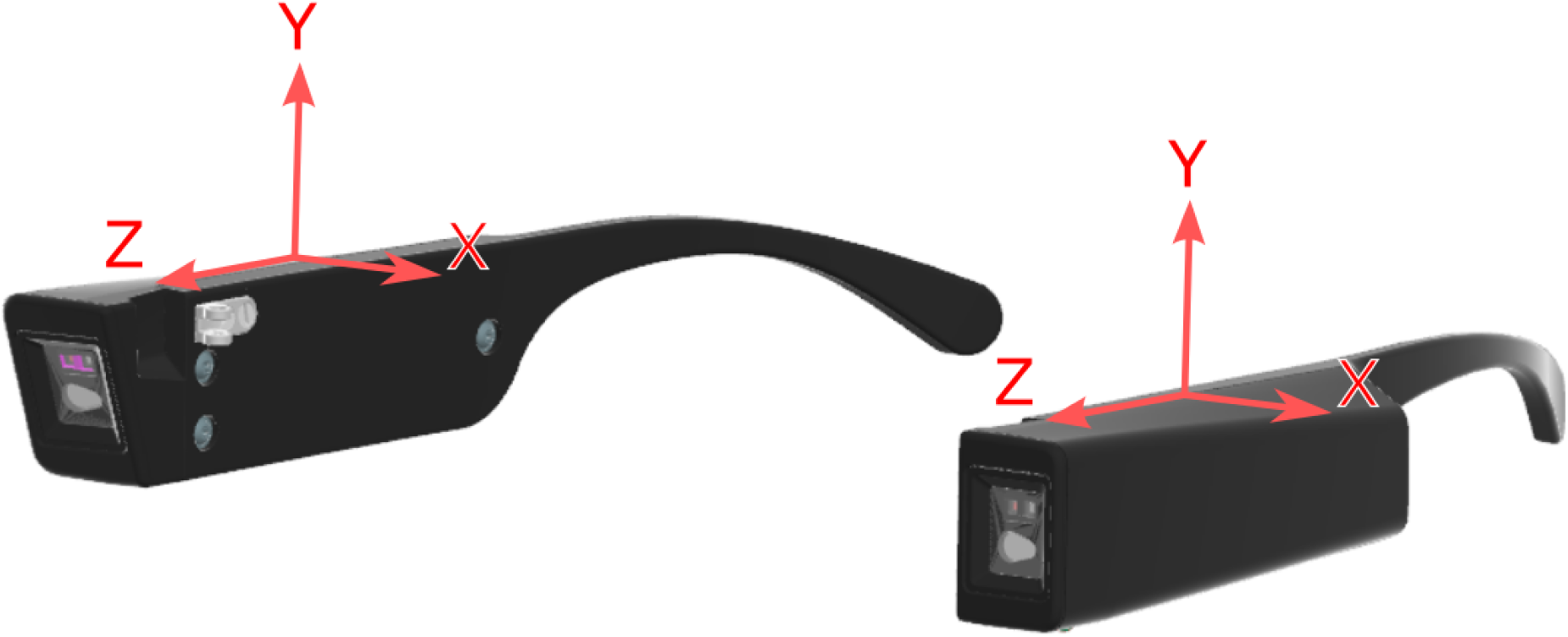
IMU Axis orientation.

### Software Interface

The open sourced VEETManager software is a convenient interface for the VEET, enabling sensor live view, device configuration, firmware and calibration updates, and full documentation.

The software has three main functions, separated into appropriate tabs for easy navigation.

● Configuration
  ○ Configure the device with sensor rates and metadata as shown in Figure 22
● Sensor live view
  ○ Individually view the 4 main sensors sampled several times per second
  ○ Individually record live view data directly to a CSV file
● Data Access
  ○ Directly navigate to a folder containing raw data

**Figure 22:**
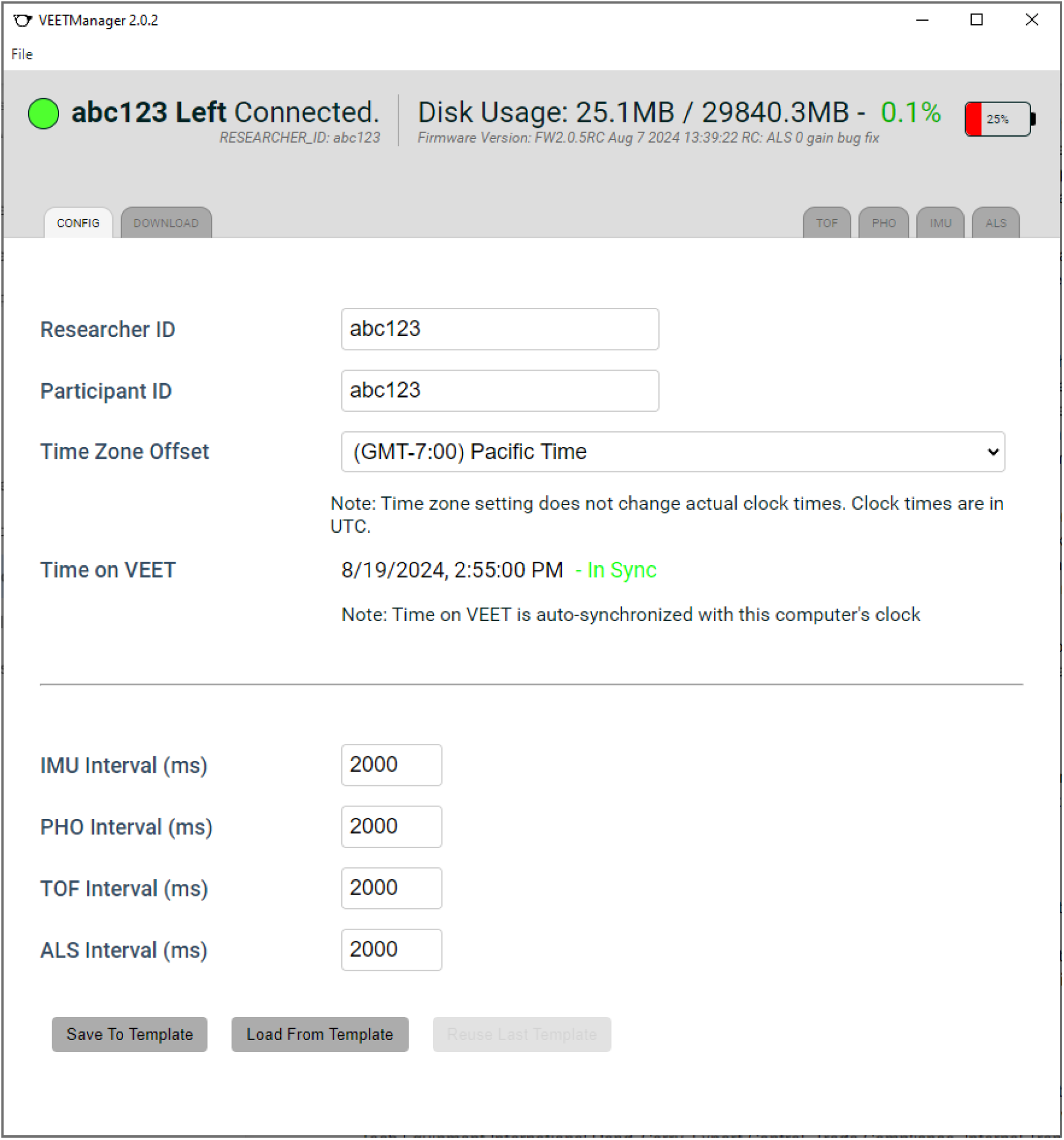
VEETManager showing active screen.

**Figure 23:**
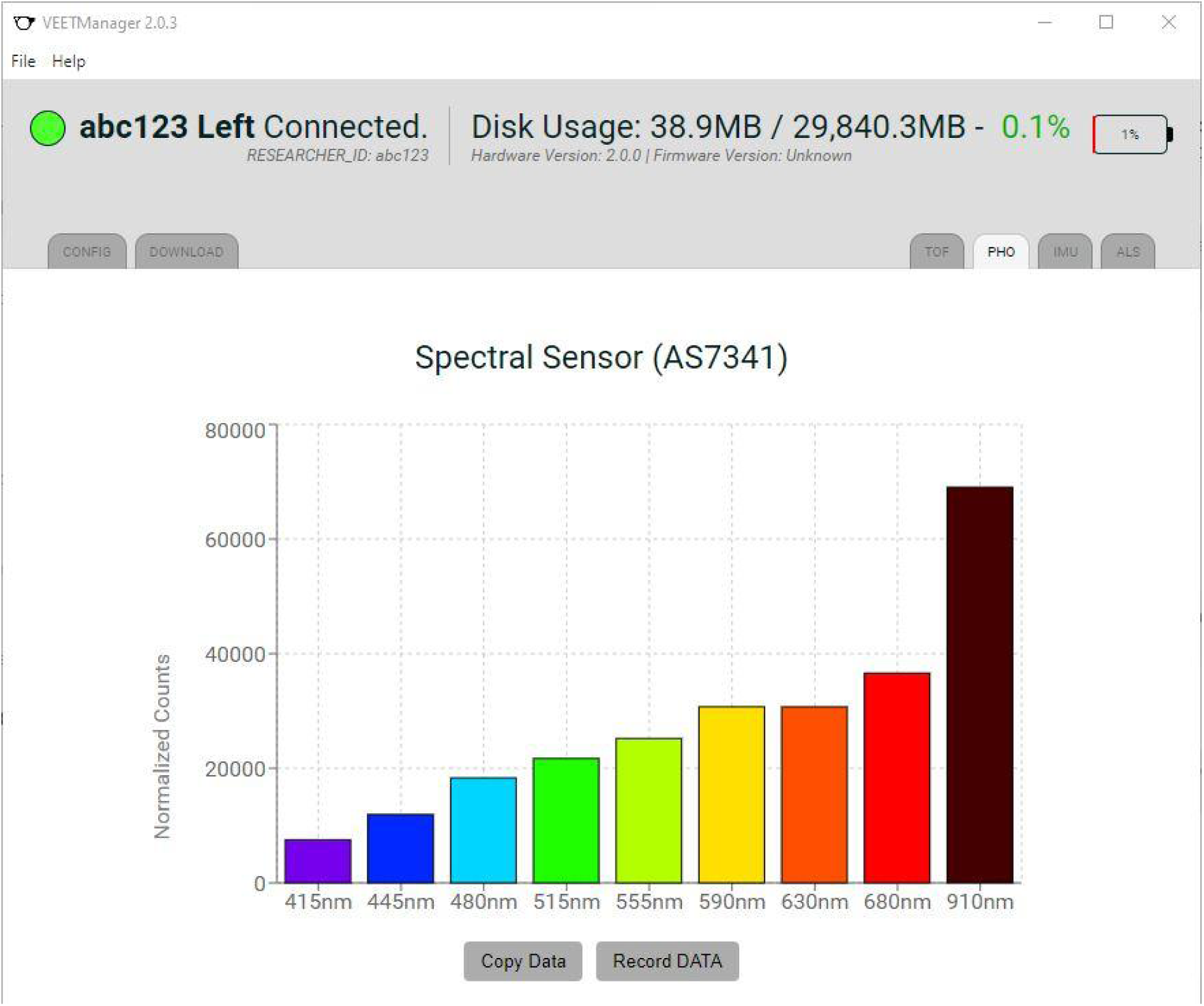
VEETManager showing typical live view screen.

Additionally, the VEETManager automatically detects firmware and software updates, prompting the user to install them. All updates are data safe and will not erase or modify any files on the VEET disk.

At this time, data visualization for logged data is not included in the software.

### Time

Time on the device is maintained in Unix time, the number of non-leap seconds that have elapsed since 00:00:00 on 1st January 1970 in Coordinated Universal Time (UTC) sometimes interchanged with the time zone Greenwich Mean Time (GMT)—and is a signed 32-bit value. Given the limited number of seconds available in this value, time on any device utilizing this method will rollover to zero and become ambiguous on January 19, 2038.

Time is maintained and recorded in whole seconds only. When the VEET is connected to the VEETManager software, time is synced automatically with the computer’s clock. During device configuration, the researcher sets the time zone offset from UTC for when and where the device is being used; this number is maintained in an information line in the data file.

The data must be post-processed to be converted to local time.

Example:

Pacific Standard Time (PST) has an offset of –8 hours from UTC.

For a Unix Epoch timestamp on device of 1704110400, this converts to the following: Date and time (UTC): Monday, January 1, 2024 12:00:00 PM (UTC-00:00)

To obtain time in the local timezone requires applying an 8 hour (28800 seconds) offset to obtain PST:

1704110400 – 28800 = 1704081600

This value converts to the following:

Date and time (Pacific Standard Time): Monday, January 1, 2024 4:00:00 AM (UTC-08:00)

Each VEET temple arm is an independent instrument that maintains its own clock. As such, exact time synchronization is not possible. There may be up to a 10-second difference in real time between temple arms after configuration due to device communication over USB.

The researcher can utilize an approach to allow closer time alignment during post-processing. Exposing the VEET to a known stimulus during a period of data collection will allow the researcher to find this event in the data file during post-processing and make time adjustments to align the data sets. The stimulus can be any specific action that will be sensed by the VEET. This method can align the time between temple arms to be within the sensing epoch configured on each device (2 seconds by default).

### Clock Drift

The microcontroller on the device utilizes a 32.768 kHz crystal oscillator as the source of a low-power mode device clock. This system has a rated tolerance of +15/-25 ppm at 35℃. This translates to –2.1 to +1.3 seconds per 24 hours. The clock may lose up to 14.7 seconds or gain 9.1 seconds from real time during the course of a week’s experiment.

### Data Access

Each VEET temple arm stores the data in a single file with a common file format known as comma-separated values (CSV). The researcher or data analyst accesses the data directly through the USB drive or through the VEETManager software. The data is recorded as time series data in a non-proprietary, fully documented format.

### Calibration

The VEET is calibrated at the time of manufacture and does not require further calibration thereafter. Nevertheless, all pertinent data is stored in the data file should the researcher consider a different calibration approach. The VEET calibration includes all light sensing ranges of the Ambient Light and Spectral Sensors. The Inertial Measurement Unit and the Time of Flight Sensors are not calibrated, but have been functionally verified as these sensors provide out-of-the-box results that are deemed accurate for this application. All methods and on board data processing will be made available.

The core components for calibration include:

Primary light source: Thorlabs SLS401 Xenon Short-Arc Light Source 240 – 2400 nm High accuracy spectrophotometer: Konica Minolta CL-500A Illuminance Spectrophotometer Optical bandpass filtering centered on each bandwidth of interest Lenses and filters to collimate and balance the light

Light tight enclosure

A reference device is measured in the light field and then compared to the VEET for calibration. Once the devices are compared, a correction factor for each channel is produced, which is loaded onto the VEET. These correction factors are loaded onto the VEET filesystem and utilized by the firmware to adjust all subsequent measurements to match the reference device.

## Conclusion

With the VEET, we are introducing a new tool that enables the ocular research community to capture egocentric light environment data to inform future models of how the eye grows and develops. The tool builds on the capabilities of previous devices and adds new modalities that we hope will fuel novel research and new insights into the factors most consequential for healthy eye development and those contributing to myopia development at population scale.

## Acknowledgements

The VEET was made possible by the team at Meta Reality Labs Research and the many academic collaborators who provided guidance, input, and feedback.

We want to especially acknowledge the invaluable early design inputs from the Ostrin Lab at the University of Houston and the important predecessor work from Bruce Weizhong Lan (ClouClip) and Lisa Ostrin (RangeLife).

